# *nGauge*: Integrated and extensible neuron morphology analysis in Python

**DOI:** 10.1101/2021.05.13.443832

**Authors:** Logan A Walker, Jennifer S Williams, Ye Li, Douglas H Roossien, Nigel S Michki, Dawen Cai

## Abstract

The study of neuron morphology requires robust and comprehensive methods to quantify the differences between neurons of different subtypes and animal species. Several software packages have been developed for the analysis of neuron tracing results stored in the standard SWC format. However, providing relatively simple quantifications and their non-extendable architecture prohibit their use for advanced data analysis and visualization. We developed *nGauge*, a Python toolkit to support the parsing and analysis of neuron morphology data. As an application programming interface (API), *nGauge* can be referenced by other popular open-source software to create custom informatics analysis pipelines and advanced visualizations. *nGauge* defines an extendable data structure that handles volumetric constructions (e.g. soma), in addition to the SWC linear reconstructions, while remaining light-weight. This greatly extends *nGauge’s* data compatibility.

## Introduction

The comparative study of neuron morphology has been a definitive aspect of contemporary neuroscience (Ramón y Cajal, 1892). Recent technological advances have enabled huge increases in the number of neuron reconstructions that can be performed in a single study into the hundreds (Gouwens et al., 2019, 2020; Jiang et al., 2021; BRAIN Initiative Cell Census Network (BICCN) et al., 2020). Multispectral labeling (Shen et al., 2020; Li et al., 2020) and large-volumetric electron microscopy (Motta et al., 2019; Yin et al., 2020; Phelps et al., 2021), in principle, allow reconstructing many more neurons within one brain or within a single common coordinate system (Wang et al., 2020). As a result, analysis techniques must be developed which allow these data to be integrated, with specific focuses on the ability to customize, automate and quickly expand processing workflows to handle large batches of individual neurons, including those reconstructed from various methods.

Neuron reconstructions are commonly abstracted as connected linear branches and stored using the SWC file format (Nanda et al., 2018). SWC files are light-weight and text-formatted that contain tab-delimited lines. Each line represents a point in the neuronal tree structure, which contains the record ID, record type (i.e., soma, axon, dendrite, etc.), X coordinate, Y coordinate, Z coordinate, the radius of the point, and the ID of the parent node to which this node links. Previously, we have also defined a volumetric SWC format where soma records are defined as a series of X-Y contour tracings along the Z axis to allow a more precise representation of soma shape (Roossien et al., 2019). Notably, the parent-child branch linkages present in SWC files result in a data model that can be understood as a directed graph, where no cycles are allowed to form. Many traditional data analysis tools do not take advantage of the underlying tree-like structure of the data, instead treating the data as a “point-cloud”.

Despite this growing need, current tools for neuron morphology calculations are largely limited to closed-form and predefined analyses, such as the popular L-Measure (Scorcioni et al., 2008) package and tools included with the various community (Peng et al., 2014; Roossien et al., 2019; Cuntz et al., 2010) or commercially-available (e.g. *Neurolucida*, MBF Biosciences; *Imaris*, Bitplane) neuron reconstruction software and plugins. Several libraries exist for the manipulation of neuron models after reconstruction, such as the TREES Toolbox (Cuntz et al., 2010) and the NeuroAnatomy Toolbox (NAT) (Bates et al., 2020), however, their APIs preclude beginner use due to their complexity. Several analysis toolkits have been introduced, such as BTMORPH (Torben-Nielsen, 2014), PyLMeasure^1^, the NAVis^2^ package, and python-Lmeasure^3^, in order to enable the quantification of neuron morphology inside of the Python programming language, which has rapidly emerged as the *lingua franca* of machine learning and data science. However, all of these tools are either limited in extensibility, or simply run other binaries in the background (which lead to large software dependencies). The recent MorphoPy (Laturnus, von Daranyi, et al., 2020; Laturnus, Kobak, et al., 2020) package solves these problems by implementing many functions in native Python code, but has only limited ability to be extended to novel metric definitions, no standardized memory structure, and no ability to produce 3D visualizations. Several software packages, such as NeuroMorphoVis (Abdellah et al., 2018) and the recent Brainrender (Claudi et al., 2020) package provide tools to prepare SWC files for complex 3D rendered figures, however, these tools are not designed to also perform quantitative analysis in the native Python environment. This data integration process for larger projects largely relies on bespoke methodologies which are limited in their reuse and accessibility.

In this report, we present *nGauge*, a software library that serves as a Python toolkit for quantifying neuron morphology. Included in the library are a collection of tools to perform standard and advanced morphometric calculations, manipulate reconstructed tree structures via SWC files, and generate visualizations within Python-native graphics libraries. We have applied *nGauge* to the analysis of several collections of published reconstructions, demonstrating the ability to build high throughput, easily understood, and reproducible bioinformatics pipelines. *nGauge* exposes a well-documented API, allowing complex morphometry analyses to be programmed quickly in conjunction with other popular bioinformatics Python software, making the library extensible and customizable to new applications. *nGauge* also operates within the Blender 3D modeling software, allowing the creation of publication-quality animations without the need for 3D rendering expertise. Finally, *nGauge* defines an extendable data structure to handle volumetric and linear neuronal constructions to greatly extend its data compatibility while remaining light-weight.

## Materials and Methods

### Library implementation

*nGauge* was implemented in Anaconda Python 3.7.6 using standard object-oriented coding practices. The results presented herein are produced using the latest version of *nGauge* as of the time of writing (0.1.2). The library makes use of other numerical methods from dependencies NumPy (Harris et al., 2020) and SciPy (Virtanen et al., 2020). Additionally, the `matplotlib` (Hunter, 2007) library is used for library plotting functions.

We implemented 103 (at time of writing) API functions which consist of single- and multivariate morphometrics, utility functions, and data structures, as described in **Results**. All implemented methods were tested with the Python `unittest` library^4^ to ensure library self-consistency. We compared the results with the output from similar functions from two previously published tools to ensure their validity (Laturnus, von Daranyi, et al., 2020; Scorcioni et al., 2008). Selected comparisons are presented in **Results**.

### Previously published data access

Previously-published neuron reconstruction data was downloaded using the bulk downloading tools on the `Neuromorpho.org` (Ascoli et al., 2007) website in SWC format (Nanda et al., 2018) from several previously-published articles (Fukunaga et al., 2012; Miyamae et al., 2017; Stokes et al., 2014). The standardized version of these SWC files was used to ensure format adherence. Additional SWC and image data were obtained from our previous study (Li et al., 2020).

### Cell type clustering

To provide a use case for how nGauge would be applied in a typical experiment, cell type clustering was performed using the above-referenced released datasets with custom python scripts. For each SWC file, the following vector of morphological parameters was calculated using nGauge: number of branch points, number of branch tips, cell dimensions, number of cell stems, average branch thickness, total path lengths, neuron volume, maximum neurite length, maximum branch order, path angle statistics, branch angle statistics, maximum branching degree, tortuosity statistics, and tree asymmetry. This collection of vectors was then used as input into the `scikit-learn` (Pedregosa et al., 2011) PCA implementation. Visual inspection of distributions was used to ensure individual clusters formed.

### Cell Mask Generation

Our cell mask generation process contains two major steps. First, a minimum convex hull of all points in the SWC file is calculated using the implemented methods in SciPy (Virtanen et al., 2020), namely the `quickhull` algorithm (Barber et al., 1996). This hull represents a 3D polygon that includes all points in the SWC file, represented as a series of lines in 3D space. After the hull is generated, the second step runs a filling. This process is applied for each neuron with a different fill value, resulting in a single-color TIFF file that can be visualized as a segmentation mask of the same size as the original image, allowing it to easily be overlaid.

### 3D Neuron visualization

All 3D visualizations were generated using the Blender 2.82.2 (Blender Foundation; blender.org) software package, following a compositing method similar to previously described (Kent, 2014). Briefly, 3D models are exported by representing each segment in an SWC file as a series of rounded cylinders, after a percentile downsampling to reduce the total number of points rendered in the 3D mesh. A decimation filter is applied to generated models to optimize the number of rendered surface points, reducing rendering time and storage requirements significantly. Standard Blender compositing techniques are then used to apply keyframes and animate scenes, as per the software documentation.

Visualizing of raw TIFF microscopy data (**Figure 7C**) was performed as follows: First, individual z-slices were exported as RGB PNG files using a script in the Fiji (Schindelin et al., 2012) image analysis software. Each slice was mapped onto the 3D model using a custom Open Shader Language (OSL) plugin (see ***Information Sharing Statement***). This allows the rendering engine to access individual z-slices without the requirement that the entire TIFF file be stored in memory.

### Performance testing

Measurement of calculation runtimes within Python was performed with the `timeit` library^5^ to run each function 4 times and automatically calculate the standard deviation using custom testing scripts (available in the ‘testing’ folder of the Github repository). L-Measure performance was measured using the Linux `time` utility to time only the compiled `lmeasure` binary, with 4 runs manually recorded from the terminal. All tests were performed on a Ubuntu Linux 20.04 server with two AMD EPYC 7351 processors, 512 GB of RAM, and all data stored on SSDs to minimize bottlenecks.

## Results

### nGauge is the center of a complete analysis environment

Neuron reconstruction experiments include three primary steps (**Figure 1A**). First, images are acquired containing the neurons of interest. Next, tracing software is used to reconstruct the neuron topology, and, finally, bioinformatic hypotheses can be tested from the resulting neuron reconstructions. These neuron reconstruction files are generally represented by the SWC format, which has been formally defined as a tabular linked list of the coordinates (Nanda et al., 2018). Because of the unique structure of this format, many general-purpose data science tools and data structures can not be efficiently applied for the analysis of neuron morphology. For this reason, we developed *nGauge* to simplify the wide variety of bioinformatics tasks, such as morphometry, model manipulation, visualization, as well as statistical analysis with the help of other python numerical libraries (**Figure 1B**).

**Figure 1:**
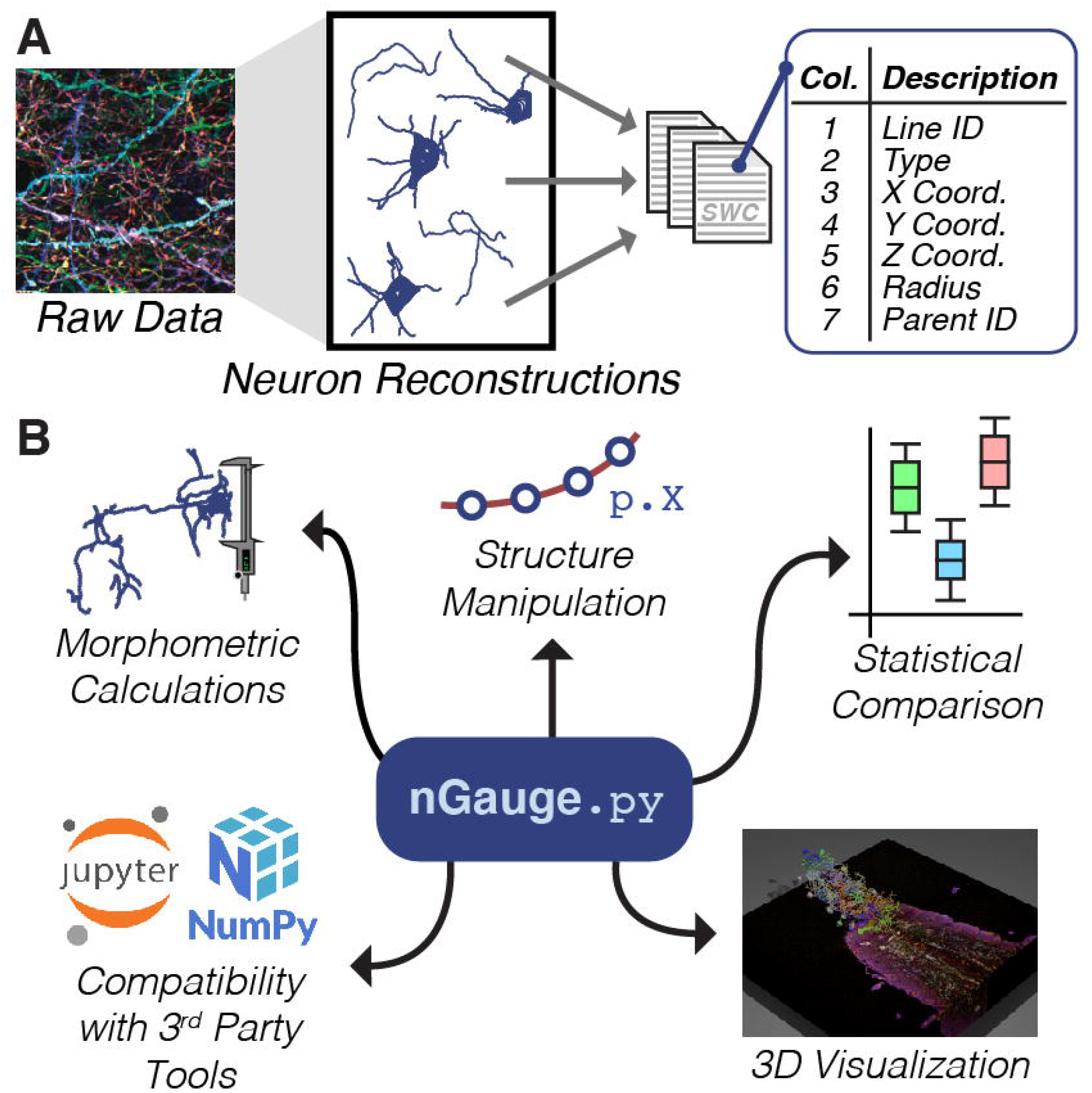
Introduction to *nGauge*. **A** Individual neuron morphologies that have been reconstructed are represented by SWC files. Each SWC file consists of a tabular list of individual points that make up the neuron tree structure; **B** The nGauge library serves as a facilitator for a variety of common Neuroinformatic tasks.

### Library structure

*nGauge* is implemented as 3 different modules, which can be installed in a single step from the Python package repository (**Figure 2A**). Each module represents an abstraction of either a single neuron, a single line in an SWC file, or the collection of utility functions used throughout the library (**Figure 2B-D**). The `Neuron` module (**Figure 2B**) stores two primary data structures. The first is a dictionary map between all soma Z-coordinates and the points which make up that “slice” of the 3D model. This data model is adapted from (Roossien et al., 2019), where the SWC format was extended to store volumetric models of somata. The second data structure stores the locations of each branch’s root node, i.e. the point at which it contacts the soma. Because each branch is a directed linked list, the only node which is needed by the Neuron module for a complete model of each of its branches is the root node of the branch. The `TracingPoint` class (**Figure 2C**) is used to represent a single SWC entry, or what would be recorded in a single line of an SWC file (**Figure 1A**), including the X, Y, and Z coordinates, as well as the point radius, and links to the `TracingPoint`’s that serve as the parent and child nodes in the linked list.

**Figure 2:**
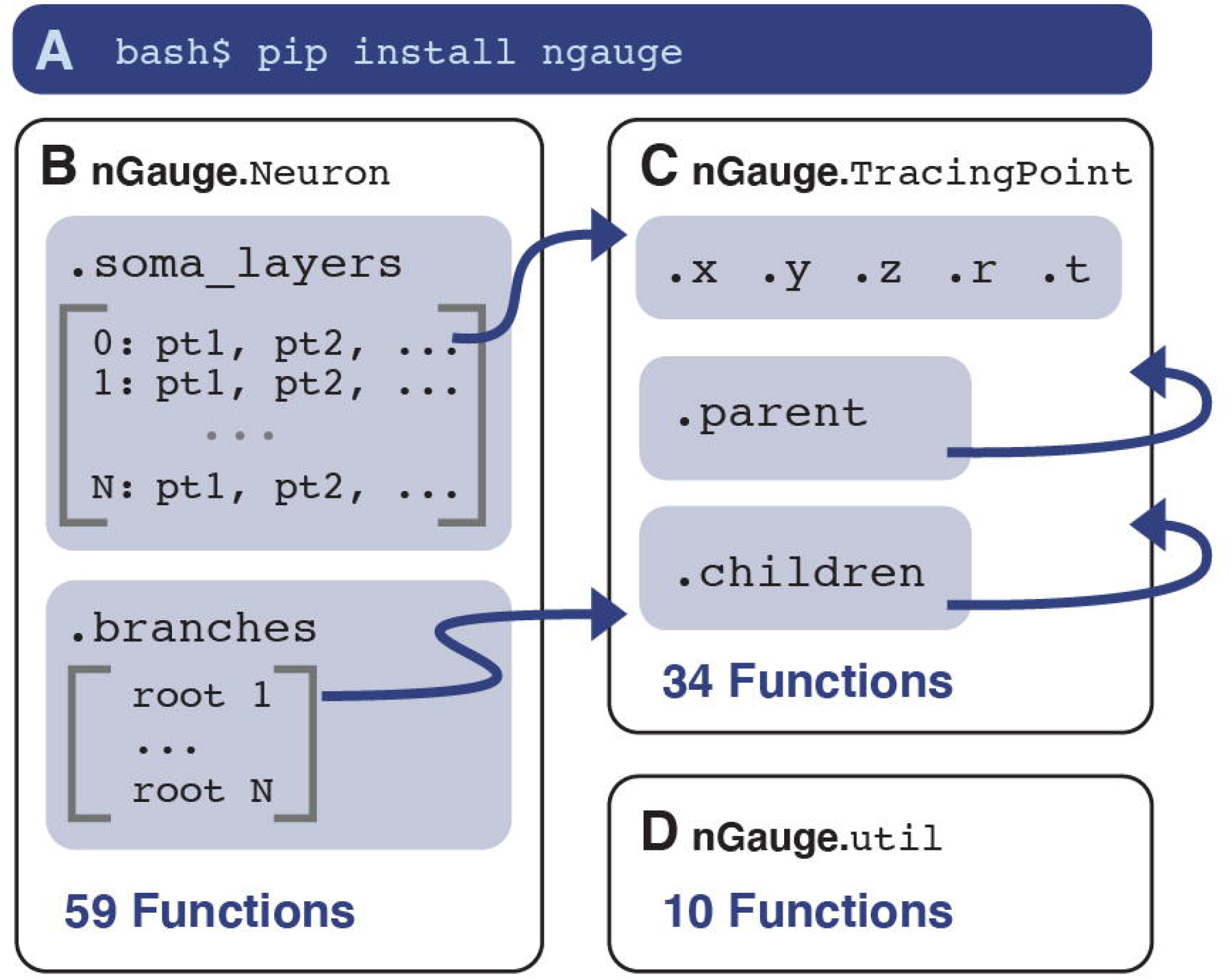
*nGauge* Library Schema. **A** nGauge is a publicly-available python library that can be installed easily in one shell command. The library is composed of 3 separate modules: Neuron (**B**), `TracingPoint` (**C**), and util (**D**). These modules implement models for an entire Neuron, a single SWC datapoint, and utility/math functions, respectively. Arrows represent cross-references between the module variables. Each module is labeled with the number of functions available at time of writing (see **Table 1** for more information).

In **Table 1**, we present a summarized list of 103 functions that are available in the current version of *nGauge*. Functions are located such that their use can match industry-standard object-oriented programming practices, leading to more readable and maintainable code. Some methods’ scope logically apply to both Neurons and TracingPoint structures (e.g., functions to calculate structure size) and are implemented in both classes.

### Introduction to nGauge usage

Care has been taken to make the use of *nGauge* as beginner-friendly as possible. To demonstrate this, we analyzed a collection of neurons from (Li et al., 2020) with our library (**Figure 3**). First, the library and data are loaded (**Figures 3A-B**). Single named morphometrics can be calculated easily by calling the methods associated with the Neuron class--in this case, the width and height of the loaded neuron (**Figure 3C**). Creating a plot of the neuron is also a single command (**Figure 3D**). While it is not shown here, plot axes and appearance parameters can be modified to get different views of the same data. Upon execution, `matplotlib` (Hunter, 2007) is loaded, allowing plots to be customized. When analyzing entire experiments or sample groups, Python list comprehension can be used to generate whole figures quickly (**Figures 3E-F**).

**Figure 3:**
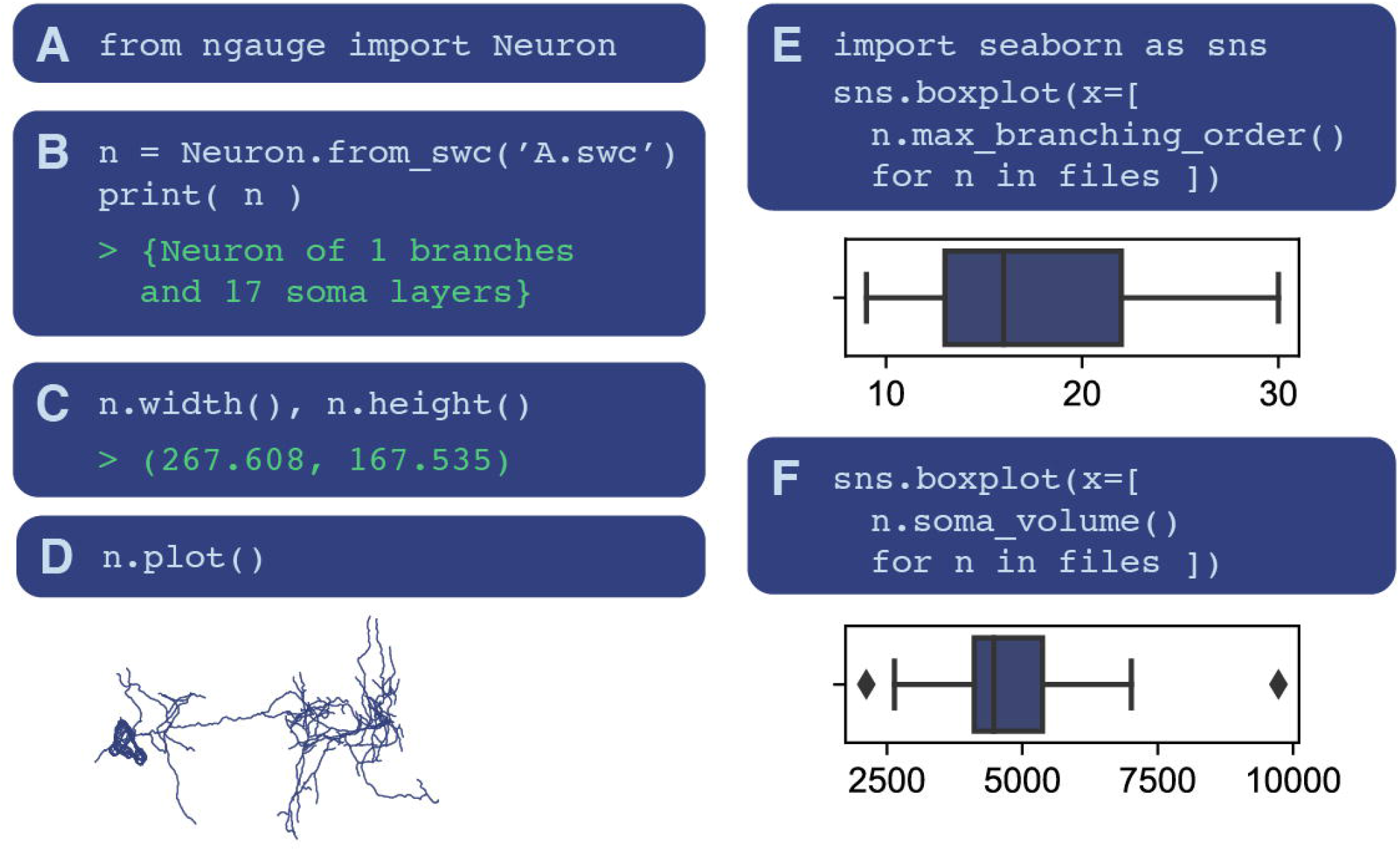
*nGauge* Usage Examples. Several code examples are included to demonstrate the processes of using nGauge. **A** Importing the library is a single command; **B** SWC files can be directly imported as Neuron objects; **C** morphometries can be easily calculated, in this case, neuron width and height (including 100% of neuron points) are calculated as members of the Neuron class; **D** Interaction with Python graphical libraries such as Matplotlib allows the generation of publication-quality figures; **E**, **F** Entire lists of files can be processed at once to run statistical analyses using python list comprehensions.

### Comparison with L-Measure

We chose to first compare our tool with L-Measure (Scorcioni et al., 2008) because it is one of the most widely-adopted and established tools for neuron morphology analysis (**Figure 4**). Additionally, several existing Python tools, such as PyLMeasure and python-Lmeasure run L-measure binaries to perform calculations in the software backend. For this comparison, we downloaded the SWC files of 42 neurons from (Stokes et al., 2014) and (Fukunaga et al., 2012), which are curated on Neuromorpho.org (Ascoli et al., 2007) (**Figure 4A, Methods**). Three representative metrics were selected to compare the tools: the number of neurite tips in the entire neuron (**Figure 4B**), the path distance of all segments of the neuron (**Figure 4C**), and the total neuron width (**Figure 4D**). As expected, the number of neurite tips and path distances are the same as calculated between the two tools (**Figures 4B-C**). The result for the total neuron width (**Figure 4D**) is more nuanced, however. The L-Measure width function is defined as the maximum distance between any two of the center 95% points along the X axis, to prevent small structures from interfering with quantification. In nGauge, this percentile becomes an option that ranges up to 100%. Finally, comparable methods between the two software packages perform faster in the *nGauge* implementation, although the speed difference varies (**Figure 4 Insets**).

**Figure 4:**
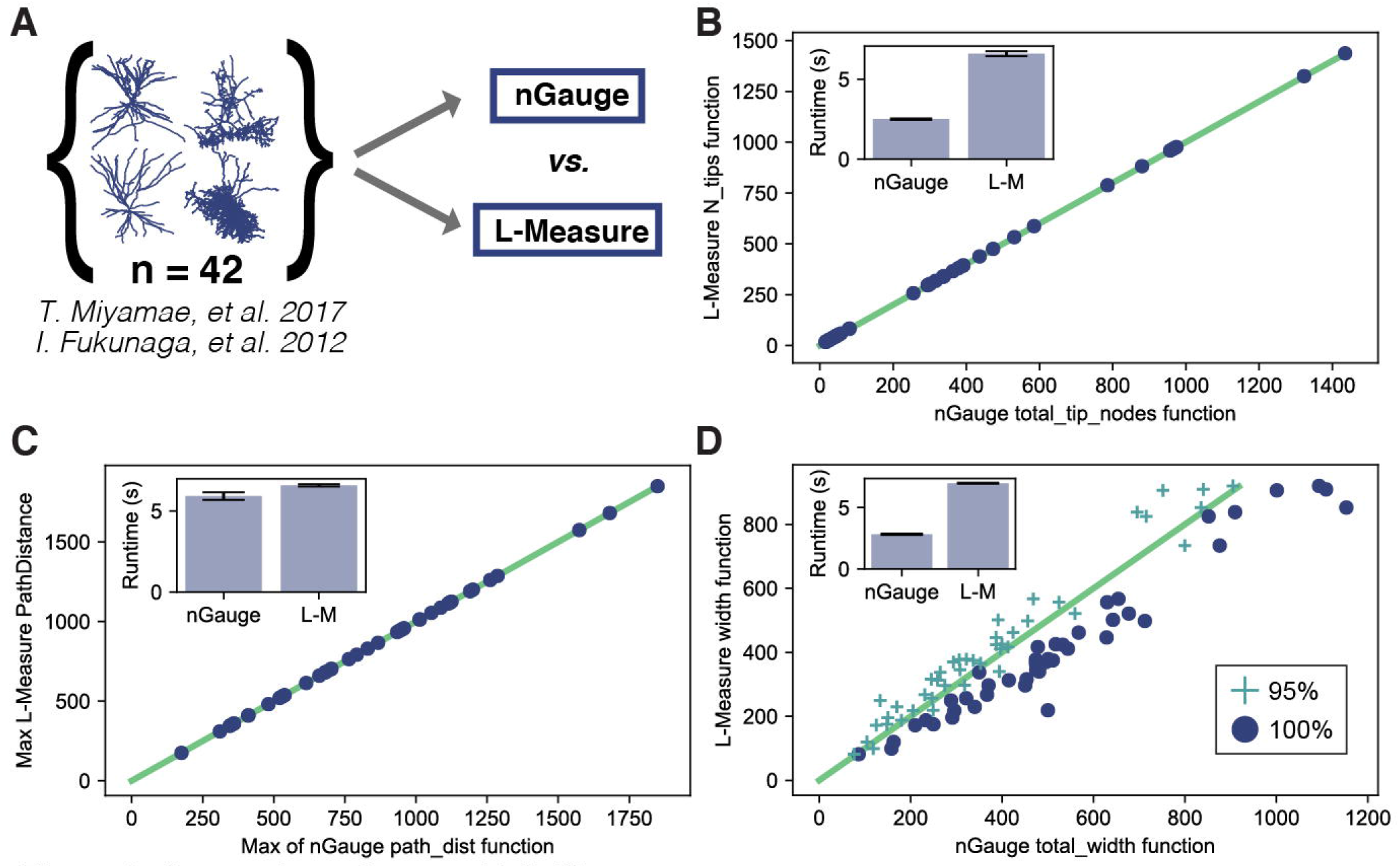
Comparing *nGauge* with L-Measure. **A** An overview of the comparison study; **B**, **C** Two example functions (tip node count and maximum path distance) produce identical output between nGauge and L-Measure; **D** Another example function (neuron width) produces similar output between nGauge and L-Measure, however, a difference of definitions produces a slight bias. Two parameter choices are shown for the nGauge result, as indicated by marker style; **Inset for each plot** nGauge scripts complete faster than their L-Measure equivalent (avg. +/- std., n = 4 per script).

### Performing advanced analysis with nGauge

In addition to quantifying basic morphometric parameters, *nGauge* includes comprehensive utility functions for advanced neuroinformatics analysis (**Table 1**). For instance, *nGauge* implements the widely used Principal Component Analysis (PCA) to identify the differences between vectors of morphometrics (Gouwens et al., 2019; Laturnus, Kobak, et al., 2020). We performed PCA on a collection of pyramidal cells and basket cells (Miyamae et al., 2017) (**Figure 5A**) and a collection of tufted cells and mitral cells (Fukunaga et al., 2012) (**Figure 5B**). Both groups are expected to form distinct morphological categories, due to their anatomical differences. We find that both comparisons yield group separation along the X-axis (principal component 1), matching what has been reported in the previous literature.

**Figure 5:**
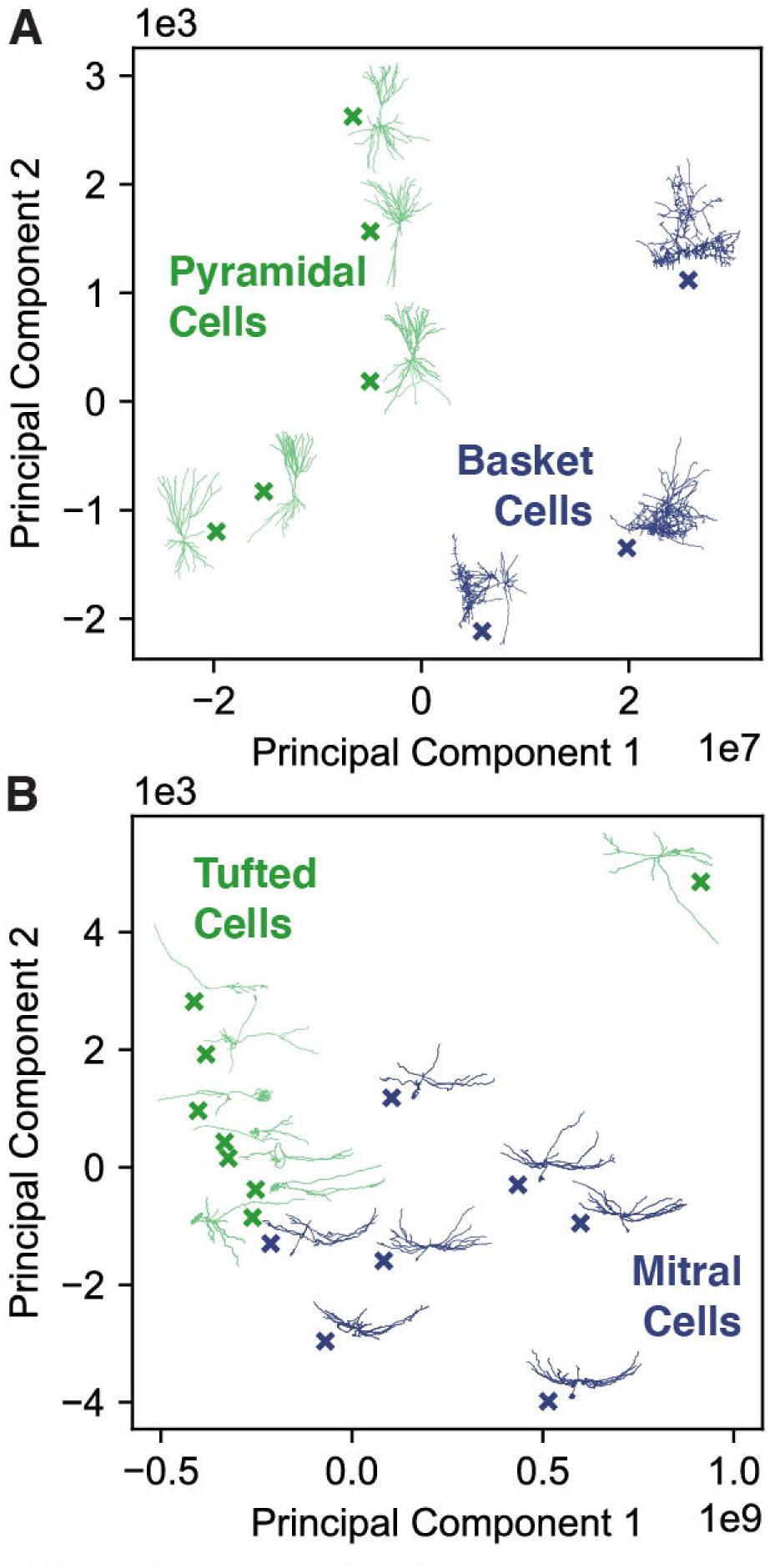
*nGauge* for Cell Type Discrimination. *nGauge* can be used to perform unbiased exploratory data analysis based on morphological parameters; **A** Pyramidal Cells are compared against Basket Cells (Miyamae et al., 2017); **B** Tufted Cells are compared against Mitral Cells (Fukunaga et al., 2012). We note that in both of these comparisons, groups form along PC1 to based on cell type. Each comparison is displayed as a principal component scatter plot and a projection of each SWC file is shown adjacent to each datapoint.

Beyond single-value morphometrics, many tools have been integrated into *nGauge* for performing advanced analysis techniques. Influenced by recent work (Laturnus, Kobak, et al., 2020), *nGauge* includes tools to calculate 2D morphometric histograms; two example cells from (Miyamae et al., 2017) are shown (**Figure 6**). These plots can serve as “fingerprints” for the morphological properties of individual neurons. In the given example the top cell is a mouse chandelier cell (NeuroMorpho ID NMO_104470) whereas the bottom cell is a Basket cell (NeuroMorpho ID NMO_104476). The difference between these two neurons can be seen in how the density of bifurcation points is much higher in the chandelier cell and also located farther from the soma (**Figure 6**, red dots).

**Figure 6:**
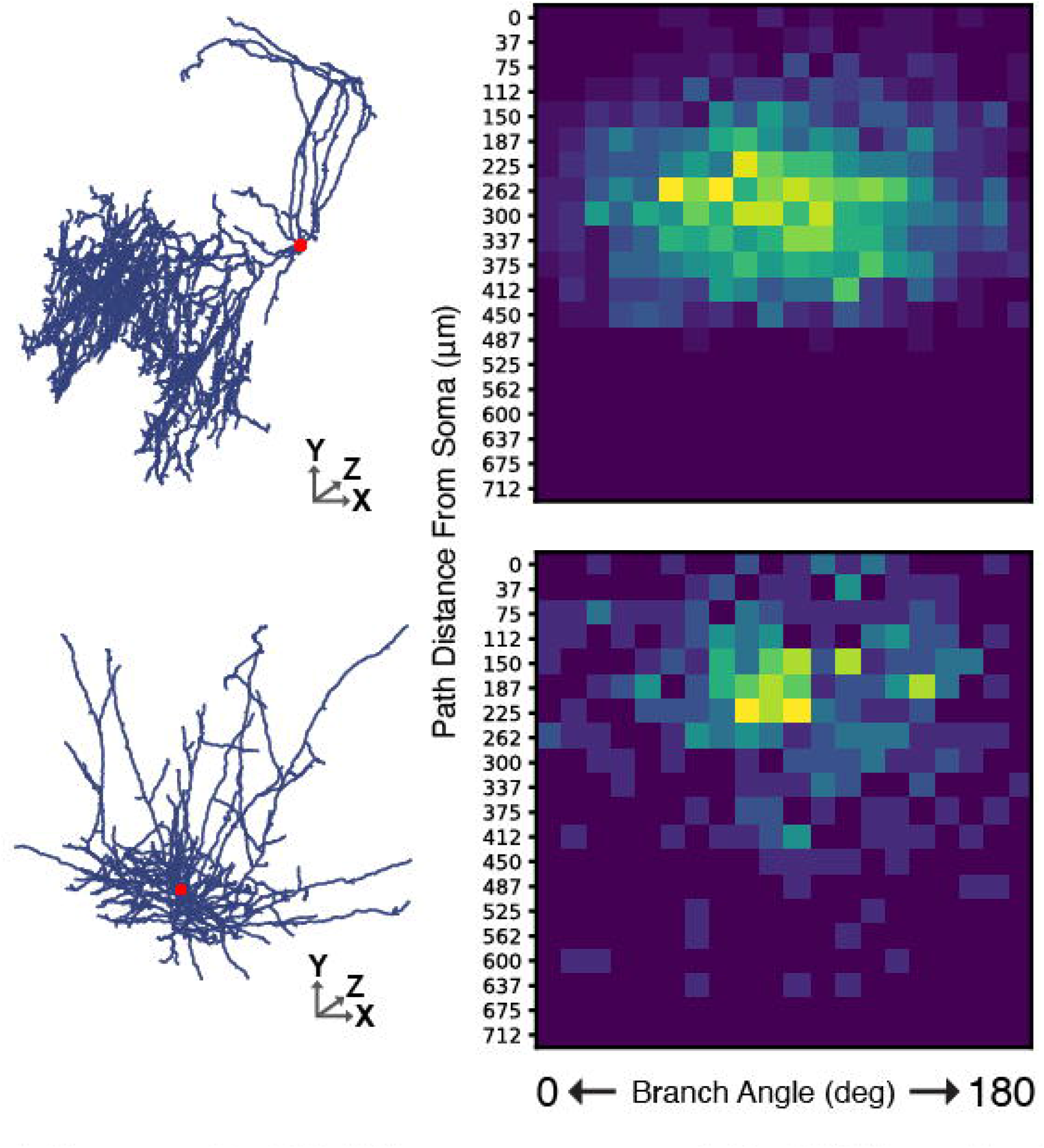
2D Histograms of Cell Morphology. Two cells (see **Results** for descriptions) are plotted as 2D histograms comparing the path distance from the soma to the branch angle for each bifurcation point in the neuron. To the left of each plot is a projection of the source SWC file, with the soma point highlighted in red.

**Figure 7:**
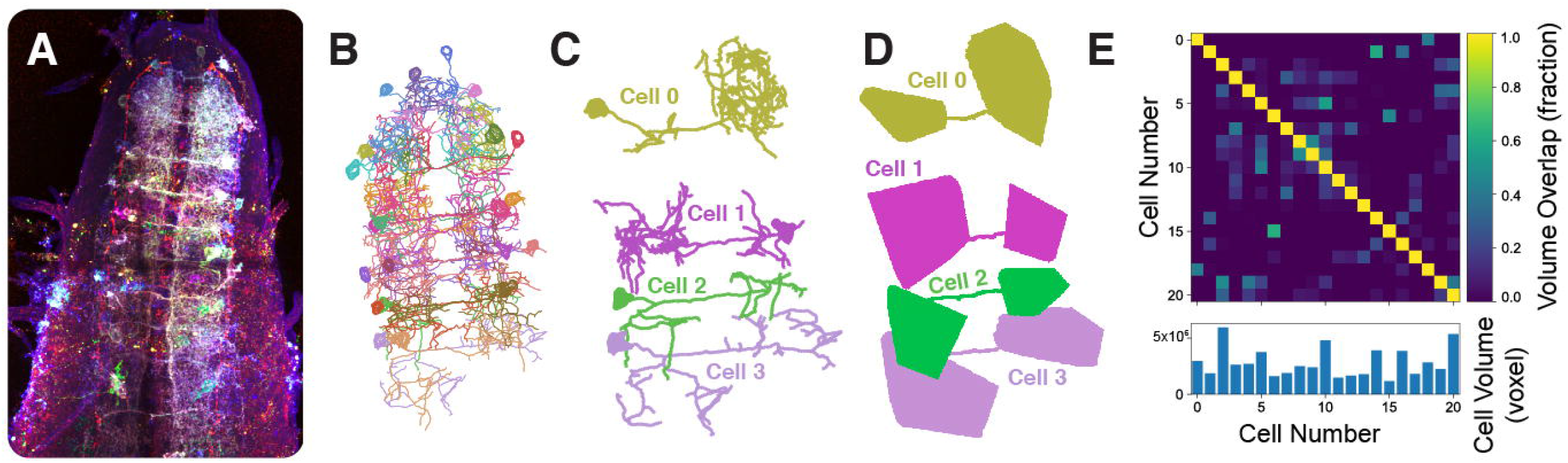
Projection Field Mapping of Multiple Neurons. We developed a novel tool for rendering the projection field volume of a specific SWC file. **A** A maximum projection of an example image from (Li et al., 2020); **B** An overview of neuron tracing reconstructed in (Li et al., 2020); **C** 4 randomly-chosen neuron reconstructions; **D** nGauge’s domain mapping tool was used to identify volumes corresponding to each SWC file in **C**; **E** A heatmap of the volume overlap percentage between each pair of samples in the experiment. Nonnegative matrix values identify cells which have overlapping domains. Each square is normalized to the volume of the cell identified in the X-axis, which is depicted in the bar plot to the bottom of the figure.

Finally, we show an example to demonstrate that nGauge can be extended to work with other Python-based scientific computation packages to create complex statistics. Using the SciPy library combined with a simple *nGauge* script, we created a unique tool for the generation of TIFF 3D masks to represent the convex hulls that enclose the extent of individual neurons. In **Figure 7**, the tool is applied to identify the neurite fields of individual Drosophila ventral nerve cord serotonergic neurons reconstructed from (Li et al., 2020). As serotonin can act as a diffusive volume transmitter (Quentin et al., 2018), each neurite field may be used to estimate the range of that serotonergic neuron’s modulation. The TIFF masks can also be used to quantify more complex geometric properties. For instance, the intersection volume between two neurons’ projection fields can be calculated using NumPy (Harris et al., 2020) as `‘np.sum(np.and(a, b))’`, or using Fiji’s `ImageCalculator` library (Schindelin et al., 2012). **Figure 7E** plots the total arborization volume of each neuron as a bar chart of total voxels (bottom) and displays this intersection volume as a heatmap between each pair of cells (top). Together, this demonstrates the utility of *nGauge* as a data structure API.

### Blender and nGauge enable advanced visualization

Visualizing neuron reconstructions in their physical context is highly valuable as it can create a direct perspective of how these neurons interact with each other and with other unreconstructed objects in the brain. We used *nGauge*’s API to create a script that renders publication-quality images and movies in the Blender 3D modeling software, which is an industry-standard open-source tool for 3D animation and visualization. We rendered the full tracing results of 182 Brainbow-labeled neurites from the CA1 region of the mouse hippocampus (Roossien et al., 2019) in two different projections (**Figures 8A-B**). These renderings, generated by only a few lines of code (available in the Github repository, **Methods**), visualize the density of the reconstruction, as well as how different somata in the reconstructed volume are positioned relative to each other. More advanced rendering techniques were used (Li et al., 2020) to visualize reconstructed serotonergic neurons of the *Drosophila* ventral nerve cord (VNC) in the context of the Bitbow fluorescence microscopy data (**Figure 8C**). Because Blender is designed for rendering still images and animations, it was possible to create a movie to display multiple angles of the neuron models (see Movie S2 in (Li et al., 2020) for example). Finally, we note that because this method is a way to convert SWC files into 3D models, it could also be used in conjunction with 3D printing technology to produce physical models of neural circuits for, e.g., educational purposes.

**Figure 8:**
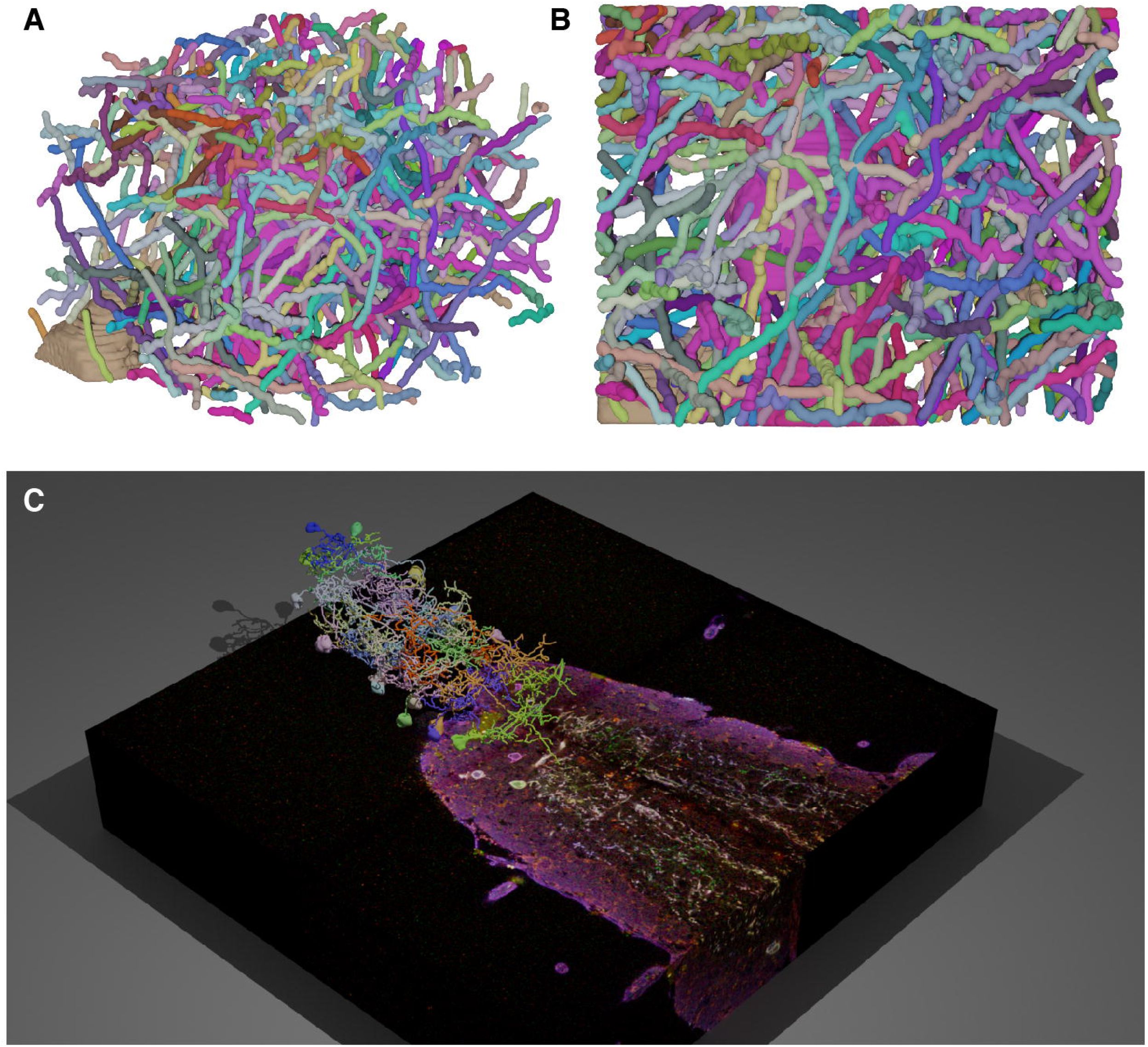
3D Modeling with nGauge and Blender. nGauge includes utilities to render publication-quality images and movies in the Blender 3D modeling software; **A, B** All somas and neurites (n=182) reconstructed in Roossien, et al. 2019 are modeled. Each panel displays a separate view of the same data; **C** Data from (Li et al., 2020) is plotted atop the raw data. A full animation of this figure is available as Movie S2 in (Li et al., 2020).

## Discussion and Conclusion

In biomedical image processing, the Fiji package (Schindelin et al., 2012) has simplified creating reproducible image processing protocols through an open environment of plugins and tools which use the Fiji data models to perform novel analyses. The development of extensible libraries such as *nGauge* are an important step to produce the same standardization in the pipelines used to analyze neuron reconstruction experiments. In its current form, *nGauge’s* library implements more than 100 morphometric calculation functions as well as provides APIs for developing new informatics tools. Notably, this simplifies the number of software tools that need to be managed and connected together to complete morphometry analysis, which lowers the learning barrier and saves time for non-informatics specialists. Combined with visualization tools, *nGauge* empowers the creation of publication-quality figures with ease. In fact, during the development of the *nGauge* project, we have already applied all of the individual modules to produce results both in publication and in preparation, finding it to be a very effective toolkit for efficient data science (Shen et al., 2020; Li et al., 2020; Dizaji et al., 2020; Duan et al., 2020).

Large-scale programs such as the NIH **B**RAIN **I**nitiative **C**ell **C**ensus **N**etwork (**BICCN**) are providing the neuroscience community with ever-expanding collections of reconstructed neuron morphology, like many other data types and modalities. Making use of this data will require a new generation of neuroinformatic data science tools that are optimized for contemporary programming techniques and are easily extensible. We believe that *nGauge* represents a significant step toward this goal, by both providing an easy way to run a large collection of “canned” analyses and by providing a platform for the experimentation and development of new metrics through a well-documented data API. As a Python library, *nGauge* can be seamlessly integrated into the most popular machine learning and data science pipelines.

In the future, we plan to continue the development of additional features for *nGauge*, such as adding tools for identifying synapse locations and performing connectivity analyses. Due to the lightweight data structure definition described here, it is straightforward to include new annotation types, such as volumetric segmentation (used in the soma here) or connectivity between tracing points in *nGauge*. We envision that *nGauge’s* open-source and expandability nature will attract contributions from the community to its public repository to make it an important toolkit of neuroscience research.

## Information Sharing Statement

*nGauge* is developed for Python 3.7 and has been tested for compatibility on the most recent version of Python at the time of writing (Python 3.9). The library is available from the Python pip package manager by executing the following command in a terminal: `‘pip install ngauge’`. The source code, documentation, and issue tracker are also available from the following Github repo: https://github.com/Cai-Lab-at-University-of-Michigan/nGauge. The provided Blender rendering tools are compatible with any version of Blender which uses a Python 3.6+ scripting interface. Installation instructions are included within the above-referenced Github repo. All data is available through the Github repository above or from the corresponding author upon reasonable request.

## Author Contributions

LAW and DC conceptualized the *nGauge* library, which was then implemented by LAW and JSW. YL and DR provided imaging and neuron reconstruction datasets which were used in library testing. LAW, JSW, YL, and NM contributed to beta testing of early versions of *nGauge* and provided comments on the library design. LAW, JSW, and DC wrote the manuscript, which was edited and approved by all authors.

## Acknowledgments

JSW received support from the University of Michigan Women in Science and Engineering Residence Program (WISE-RP) Judith Cram Memorial Fund Research Award. LAW and DC received support from NSF-1707316 (Neuronex-MINT), NIH-RF1MH123402, and NIH-RF1MH124611. The authors thank Fred Shen for his comments on an early version of the library. LAW thanks Chris Midkiff for his comments on figure design and example code clarity.

## Figure and Table Captions

**<u>Figure 1</u> Introduction to *nGauge***

**A** Individual neuron morphologies that have been reconstructed are represented by SWC files. Each SWC file consists of a tabular list of individual points that make up the neuron tree structure; **B** The *nGauge* library serves as a facilitator for a variety of common Neuroinformatic tasks.

**<u>Figure 2</u> *nGauge* Library Schema**

**A** *nGauge* is a publicly-available python library that can be installed easily in one shell command. The library is composed of 3 separate modules: `Neuron` (**B**), `TracingPoint` (**C**), and `util` (**D**). These modules implement models for an entire Neuron, a single SWC datapoint, and utility/math functions, respectively. Arrows represent cross-references between the module variables. Each module is labeled with the number of functions available at time of writing (see **Table 1** for more information).

**<u>Figure 3</u> *nGauge* Usage Examples**

Several code examples are included to demonstrate the processes of using *nGauge*. **A** Importing the library is a single command; **B** SWC files can be directly imported as Neuron objects; **C** morphometrics can be easily calculated, in this case, neuron width and height (including 100% of neuron points) are calculated as members of the Neuron class; **D** Interaction with Python graphical libraries such as Matplotlib allows the generation of publication-quality figures; **E, F** Entire lists of files can be processed at once to run statistical analyses using python list comprehensions.

**<u>Figure 4</u> Comparing *nGauge* with L-Measure**

**A** An overview of the comparison study; **B, C** Two example functions (tip node count and maximum path distance) produce identical output between *nGauge* and L-Measure; **D** Another example function (neuron width) produces similar output between *nGauge* and L-Measure, however, a difference of definitions produces a slight bias. Two parameter choices are shown for the nGauge result, as indicated by marker style; **Inset for each plot** *nGauge* scripts complete faster than their L-Measure equivalent (avg. +/- std., n = 4 per script).

**<u>Figure 5</u> *nGauge* for Cell Type Discrimination**

*nGauge* can be used to perform unbiased exploratory data analysis based on morphological parameters; **A** Pyramidal Cells are compared against Basket Cells (Miyamae et al., 2017); **B** Tufted Cells are compared against Mitral Cells (Fukunaga et al., 2012). We note that in both of these comparisons, groups form along PC1 based on cell type. Each comparison is displayed as a principal component scatter plot and a projection of each SWC file is shown adjacent to each datapoint.

**<u>Figure 6</u> 2D Histograms of Cell Morphology**

Two cells (see **Results** for descriptions) are plotted as 2D histograms comparing the path distance from the soma to the branch angle for each bifurcation point in the neuron. To the left of each plot is a projection of the source SWC file, with the soma point highlighted in red.

**<u>Figure 7</u> Projection Field Mapping of Multiple Neurons**

We developed a novel tool for rendering the projection field volume of a specific SWC file. **A** A maximum projection of an example image from (Li et al., 2020); **B** An overview of neuron tracing reconstructed in (Li et al., 2020); **C** 4 randomly-chosen neuron reconstructions; **D** *nGauge’s* domain mapping tool was used to identify volumes corresponding to each SWC file in **C**; **E** A heatmap of the volume overlap percentage between each pair of samples in the experiment. Nonnegative matrix values identify cells which have overlapping domains. Each square is normalized to the volume of the cell identified in the X-axis, which is depicted in the bar plot to the bottom of the figure.

**<u>Figure 8</u> 3D Modeling with *nGauge* and Blender**

*nGauge* includes utilities to render publication-quality images and movies in the Blender 3D modeling software; **A, B** All somas and neurites (n=182) reconstructed in Roossien, et al. 2019 are modeled. Each panel displays a separate view of the same data; **C** Data from (Li et al., 2020) is plotted atop the raw data. A full animation of this figure is available as Movie S2 in (Li et al., 2020).

**<u>Table 1</u> Implemented *nGauge* Functions**

All functions implemented in *nGauge*. Along the left hand side, functions are divided into different modules and function types.

## Footnotes

1 https://pypi.org/project/pylmeasure/

2 https://navis.readthedocs.io/en/latest/source/other_libraries.html

3 https://github.com/ajkswamy/python-Lmeasure

4 https://docs.python.org/3/library/unittest.html

5 https://docs.python.org/3/library/timeit.html

