## Supplementary material for "*nGauge*: Integrated and extensible neuron morphology analysis in Python": Table 1

| | Type | Function | Description | Included in LMeasure |
|---|---|---|---|---|
| Neuron | Structure | *Add Branch* | Appends a TracingPoint object as a new branch root to the neuron | NA |
| | | *Center Soma* | Translates all Neuron coordinates such that the soma centroid falls on the origin (0, 0, 0). | NA |
| | | *Scale* | Scales the neuron by a given factor along each axis | NA |
| | | *Translate* | Transforms all coordinates along a given distance along each axis | NA |
| | | *Rotate* | Rotates all coordinates by given angles around each axis | NA |
| | | *Add Soma Points* | Appends a list of points as a new soma layer to the neuron | NA |
| | Morphometric | *Width* | Calculates the width of the bounding box around a given neuron | Yes |
| | | *Height* | Calculates the height of the bounding box around a given neuron | Yes |
| | | *Depth* | Calculates the depth of the bounding box around a given neuron | Yes |
| | | *Soma Centroid* | Calculates the center-of-mass of all the soma points | No |
| | | *Total Root Branches* | Calculates the number of branch roots included in the neuron | Yes |
| | | *Total Child Nodes* | Calculates the number of child nodes included in the neuron | Yes |
| | | *Total Tip Nodes* | Calculates the total tip (end of branch) nodes included in the neuron | Yes |
| | | *Total Bifurcation Nodes* | Calculates the total number of bifurcation (branching point) nodes included in the neuron | Yes |
| | | *Soma Volume* | Calculates the total volume of the neuron's soma points | Yes |
| | | *Soma Surface Area* | Calculates the total surface area of the neuron's soma points | Yes |
| | | *Soma Slice Perimeter* | Calculates the perimeter of each Z-slice defined for the soma | No |
| | | *Persistence Diagram* | Runs a given function over all segments and returns a list of results, can be used to implement persistence diagrams | No |
| | | *Arborization Distance* | Calculates the distance from the soma centroid to the average of all branch coordinates | No |
| | | *Maximum Branch Angle* | Finds the largest branching angle in the Neuron | Yes |
| | | *Minimum Branch Angle* | Finds the minimum branching angle in the Neuron | Yes |
| | | *Average Branch Angle* | Calculates the average of all branching angles in the neuron | Yes |
| | | *Maximum Path Angle* | Finds the largest path angle in the Neuron | Yes |
| | | *Median Path Angle* | Finds the median path angle in the Neuron | Yes |
| | | *All Branch Angles* | Returns a list of all branch angles in the neuron | Yes |
| | | *Max Tortuosity* | Finds the largest tortuosity in the Neuron | No |
| | | *Median Tortuosity* | Finds the median tortuosity in the Neuron | No |
| | | *Maximum Branching Order* | Calculates the highest branching order achieved by any tip node | Yes |
| | | *All Segment Lengths* | Returns a list of all segment lengths in the Neuron | No |
| | | *All Path Angles* | Returns a list of all path angles in the Neuron | No |
| | Distributions | *Branch Order Counts* | Returns a count list of all branch orders | No |
| | | *Branch Order Histogram* | Returns a histogram of all branch orders of all bifurcations | No |
| | | *Path Angle Histogram* | Returns a histogram of all path angles | No |
| | | *Path Distance to Soma Histogram* | Returns a histogram of all path distances to the soma | No |

| | | | | |
|---|---|---|---|---|
| | | *Euclidean Distance to Soma Histogram* | Returns a histogram of Euclidean distances from each TracingPoint to the soma | No |
| | | *Thickness Histogram* | Returns a histogram of all TracingPoint thicknesses | No |
| | | *Branch Angle Histogram* | Returns a histogram of all branch angles | No |
| | | *Path Angles Histogram* | Returns a histogram of all path angles | No |
| | | *Branch Angles by Branch Orders* | Performs a 2D histogram of branch angles and branch orders | No |
| | | *Branch Angles by Path Distance* | Performs a 2D histogram of branch angles and path distances | No |
| | | *Thickness by Branch Order* | Performs a 2D histogram of thickness by branch order | No |
| | | *Thickness by Path Distance* | Performs a 2D histogram of thickness by path distance | No |
| | **Utility** | *Get Main Branch* | Finds the longest branch in the neuron and returns it | NA |
| | | *Plot* | Generates a X, Y, or Z maximum projection of an entire neuron | No |
| | | *Iterate All Points* | Iterates through all TracingPoint objects as a Python generator object | NA |
| | | *From SWC* | Imports a SWC file from either a file on the disk or a Python string object | NA |
| | | *To SWC* | Exports the neuron as a SWC to either a file or Python string object | NA |
| | | *Fix Parents* | Ensures that all nodes in the Neuron are correctly linked in the tree structure | NA |
| **TracingPoint** | **Structure** | *Add Child* | Adds another TracingPoint as either an extension of a single branch if one does not already exist, or as a second sub-branch | NA |
| | | *Get Tip Nodes* | Returns a list of all tip node TracingPoints | Yes |
| | | *Get Bifurcation Nodes* | Returns a list of all bifurcation node TracingPoints | Yes |
| | | *Get All Nodes* | Returns a list of all node TracingPoints | NA |
| | | *Get All Segments* | Returns a list of all TracingPoint pairs representing individual segments along the branch | NA |
| | | *Path Distance to Child* | Returns the path distance from the given point and a given child node | Yes |
| | | *Next Bifurcation Point* | Finds the nearest neighbor bifurcation point in the branch tree | NA |
| | | *Select Nodes* | Returns all nodes which return true for a given test function, used to select TracingPoints which meet a given criteria | NA |
| | **Morphometric** | *Total Children* | Counts the number of subbranches present at this TracingPoint | Yes |
| | | *Total Child Nodes* | Recursively counts the number of TracingPoints that make up all subbranches of this TracingPoint | Yes |
| | | *Total Tip Nodes* | Counts the total number of child nodes which are tips nodes | Yes |
| | | *Total Bifurcation Nodes* | Counts the total number of child nodes which are bifurcation nodes | Yes |
| | | *Is Tip Node* | Returns True if the node is a tip node | Yes |
| | | *Is Bifurcation Node* | Returns True if the node is a bifurcation node | Yes |
| | | *Is Root Node* | Returns True if the node is a root node | Yes |
| | | *Path Distance to Root* | Calculates the total path distance from this node to the soma | Yes |
| | | *Width* | Calculates the width of a bounding box around the branch | Yes |
| | | *Height* | Calculates the height of a bounding box around the branch | Yes |
| | | *Depth* | Calculates the depth of a bounding box around the branch | Yes |
| | | *Volume* | Calculates the volume of a bounding box around the branch | Yes |

| | | | | |
|---|---|---|---|---|
| | | *Distance to Ends* | Calculates the Euclidean distance to each child tip node | Yes |
| | | *Path Distance to Ends* | Calculates the path distance to each child tip node | Yes |
| | | *Partition Asymmetry* | Calculates the partition asymmetry of this node | Yes |
| | | *Neurite Tortuosity* | Calculates the tortuosity of the neurite | No |
| | | *Branching Order* | Calculates the branching order of this node | Yes |
| | **Utility** | *Euclidean Distance* | Calculates the Euclidean distance between two TracingPoints | NA |
| | | *Slice Surface Area* | Calculates the surface area of a collections of points that define a slice | NA |
| | | *Slice Perimeter* | Calculates the perimeter of a collection of points that define a slice | NA |
| | | *To SWC* | Exports this branch to a SWC file | NA |
| | | *Fix Parents* | Corrects linkages for all TracingPoint children of this node | NA |
| **Utility** | **Math** | *Rotation Matrix* | Calculates a standard rotation matrix of three angles | NA |
| | | *Tangent from Points* | Calculates a tangent vector from a list of points | NA |
| | | *Dot Product* | Performs the dot product of two vectors | NA |
| | | *Absolute Dot Product* | Calculates the absolute value of the dot product | NA |

**Table 1 Implemented *nGauge* Functions**
All functions implemented in *nGauge*. Along the left hand side, functions are divided into different modules and function types.
